# Fibronectin orchestrates extracellular matrix composition and cardiac outflow tract elongation in *Xenopus laevis*

**DOI:** 10.64898/2026.03.18.712624

**Authors:** Javiera Jorquera, Isidora Sovino, Catalina Jara-Gonzalez, Ignacio Rosales, Paula G Slater, Cecilia Arriagada

**Affiliations:** Centro de Biología Celular y Biomedicina (CEBICEM), Facultad de Ciencias, Universidad San Sebastián, Avenida del Valle Norte 725, Huechuraba, Santiago, Chile. 8580702; Departamento de Ciencias Biológicas y Químicas, Facultad de Ciencias, Universidad San Sebastián, Campus Los Leones, Lota 2465, Providencia, Santiago, Chile; Laboratorio de Neuro-regeneración y metabolismo, Fundación Ciencia & Vida. Avenida del Valle Norte 725, Huechuraba, Santiago, Chile. 8580702

**Keywords:** Fibronectin Heart Development, Extracellular Matrix, Xenopus

## Abstract

Congenital heart defects frequently arise from alterations in the elongation of the cardiac outflow tract (OFT). Proper elongation of the OFT depends on the coordinated deployment of progenitor cells from the second heart field (SHF) and on dynamic interactions with the extracellular matrix (ECM). Among ECM components, fibronectin (Fn1) and tenascin-C (TnC) have emerged as key regulators of cardiac morphogenesis. Studies in mouse embryos have shown that mesodermal Fn1 is required to maintain proper TnC localization within SHF cells. To study heart development, mammalian models are challenging to use because of their in utero development. This limitation highlights the need for alternative models with external development, where direct observation is possible; however, in these systems, the cellular organization of the SHF and the dynamics of its ECM environment remain poorly characterized

Here, we investigated the cellular and extracellular architecture of SHF cells localized to the dorsal pericardial wall (DPW) during heart development in *Xenopus laevis.* We show that SHF cells undergo a stage-dependent transition from a predominantly monolayered organization at NF35 to a multilayered structure at NF42. This transition is accompanied by dynamic remodeling of the ECM, characterized by increased expression of Fn1, TnC, and Collagen I (ColI) and by redistribution of ECM components within the DPW.

Functional experiments revealed that depletion of Fn1 disrupts cardiac morphogenesis, leading to shortening of the OFT and reduced ventricular size. Moreover, loss of Fn1 decreases TnC and ColI levels and alters the spatial organization of TnC within the DPW, indicating that Fn1 is required for proper ECM assembly within the SHF cells.

These findings identify Fn1 as a key regulator of ECM assembly within the DPW and highlight how ECM remodeling contributes to the organization of SHF progenitor cells during OFT elongation. Altogether, we demonstrated that *Xenopus laevis* is a powerful model for studying ECM-driven mechanisms of cardiac morphogenesis.

## Introduction

Congenital heart defects are among the most common developmental disorders in humans and, in many cases, arise from early alterations in the morphogenetic processes that shape the embryonic heart, particularly the elongation of the outflow tract (OFT) (Neeb et al., 2013; Tsao et al., 2022) and the formation of the cardiac chambers (Bruneau, 2008). Proper elongation of the OFT depends on the precise coordination of proliferation, migration, and organization of cardiac progenitor populations, as well as on dynamic interactions between cells and the surrounding extracellular matrix (ECM) (Buckingham et al., 2005; Silva et al., 2021)

One of the principal cellular lineages contributing to OFT elongation is the second heart field (SHF), a population of multipotent progenitor cells located adjacent to the developing heart that are progressively incorporated into the arterial and venous poles of the heart tube (Buckingham et al., 2005; Kelly et al., 2014). The ECM is a fundamental regulator of embryonic morphogenesis, as it not only provides structural support but also delivers biochemical and mechanical cues that influence cell polarity, migration, cell shape, and cell fate decisions (Hynes, 2009; Rozario and DeSimone, 2010). During cardiac development, ECM components are dynamically deposited and reorganized around cardiac progenitors, contributing to tissue integrity and to the transmission of forces necessary for cardiac tube elongation (Le A. Trinh and Stainier, 2004; Czirok et al., 2006). Despite important advances in identifying genetic regulators of cardiogenesis, the mechanisms by which the ECM contributes to tissue organization and cardiac progenitor behavior remain poorly unknown. Several ECM proteins, including fibronectin (Fn1), tenascin-C (TnC), collagen type I (Col I), and laminin, have been described as key regulators of heart formation (Silva et al., 2021). Among these, Fn1 and TnC have emerged as important regulators of cardiac morphogenesis. Fn1 is a large glycoprotein that assembles into fibrillar networks through integrin-dependent interactions and plays a central role in cell–ECM adhesion and in the organization of the tissue microenvironment (Hynes, 1986; Pankov and Yamada, 2002). Genetic studies in different vertebrate models have demonstrated that Fn1 is essential for developmental processes that require extensive tissue remodeling, such as gastrulation, somitogenesis, and cardiovascular organogenesis (Astrof, Crowley, et al., 2007; Astrof, Kirby, et al., 2007,; Chen et al., 2015; George et al., 1993; Le A. Trinh & Stainier, 2004; Warkala et al., 2021)

In contrast, TnC is a hexameric ECM protein that is highly expressed during embryonic development and re-expressed in pathological contexts, such as tissue repair and cancer (Chiquet-Ehrismann et al., 1986; Chiquet-Ehrismann and Chiquet, 2003; Midwood et al., 2016). Unlike Fn1, TnC has been described as an anti-adhesive or adhesion-modulating molecule that regulates cell shape, tissue plasticity, and integrin-mediated signaling (Radwanska et al., 2017). During cardiac development, TnC is preferentially localized in regions undergoing intense morphogenetic remodeling, including the OFT and domains associated with the SHF (Imanaka-Yoshida et al., 2003; Imanaka-Yoshida and Aoki, 2014). Accumulating evidence indicates that Fn1 and TnC functionally interact within the ECM. *In vitro* studies have shown that TnC can associate with Fn1 fibrils and modulate integrin-mediated adhesion and signaling, thereby influencing the balance between cell adhesion and tissue plasticity (Chiquet-Ehrismann et al., 1988; Orend et al., 2003; To and Midwood, 2010; Midwood et al., 2016). In the SHF of *Mus musculus* (mouse) embryos, loss of mesodermal Fn1 leads to mislocalization of TnC and disruption of ECM organization, which is associated with defects in OFT elongation and cardiac morphogenesis (Arriagada et al., 2025). These findings suggest that a finely regulated balance between adhesive and modulatory ECM components is crucial for maintaining SHF tissue architecture.

Studying cardiac development in mouse models can be technically challenging because it occurs in utero. Therefore, comparative studies across vertebrate models, particularly those with external embryonic development, are valuable for advancing our understanding of cardiac morphogenesis and for distinguishing conserved mechanisms from lineage-specific processes. *Xenopus laevis* has been widely used as a model organism for studying early vertebrate development due to its external development and ease of experimental manipulation. Moreover, *Xenopus* embryogenesis proceeds rapidly, enabling the study of the entire cardiovascular developmental process within a relatively short time frame (Kaltenbrun et al., 2011; Carotenuto et al., 2023). Although the *Xenopus* heart displays a simpler anatomy than that of mammals, the cardiogenic signaling pathways as well as the fundamental processes of cardiac tube formation, OFT elongation, and SHF progenitor contribution to OFT growth are conserved (Sater and Jacobson, 1989; Kolker et al., 2000; Mohun et al., 2000; Schneider and Mercola, 2001; Brade et al., 2007; Lee and Saint-Jeannet, 2011). However, the organization of the SHF and the composition and dynamics of its ECM in *Xenopus* remain relatively poorly characterized.

In this study, we investigated the cellular and extracellular architecture of the SHF associated with the dorsal pericardial wall (DPW) during cardiac development in *Xenopus laevis*, with a particular focus on dynamic ECM remodeling. We characterized tissue organization, cell morphology, and epithelial features of SHF cells in the DPW at different developmental stages and analyzed the temporal and spatial distribution of key ECM components, including Fn1, TnC, ColI, and laminin. In addition, we evaluated the functional role of Fn1 in OFT elongation and ECM organization, as well as its impact on TnC localization.

Together, our results provide new insights into how ECM remodeling contributes to SHF architecture and cardiac morphogenesis in a non-amniote vertebrate and reveal both conserved and divergent aspects of Fn1–TnC interactions during heart development. These findings position *Xenopus laevis* as a valuable comparative model for understanding ECM-dependent mechanisms relevant to the origin of congenital heart defects.

## Materials and Methods

### *Xenopus laevis* embryo breeding and maintenance

Adult female and male *Xenopus laevis* were pre-injected with 50 μL human chorionic gonadotropin (hCG, 1000 IU/mL), 72 h later they received a booster injection of 700 μL and 300 μL hCG (1000 IU/mL), respectively. Females were crossed with adult males by amplexus, and external fertilization occurred approximately 12 h post-injection. Fertilized eggs were collected and dejellied by incubation in 2 % cysteine solution for 5 min, followed by washes with distilled water and 0.1X Barth solution, as previously described (Slater and Larraín, 2021). Embryos were maintained in sterilized glass dishes at 23°C until reaching the desired Nieuwkoop and Faber (NF) developmental stages (Zahn et al., 2022). All animal procedures were approved by the Bioethical and Biosafety Committee of Universidad San Sebastián.

### Whole-mount immunofluorescence

Embryos at NF35 and NF42 stages were fixed in 4% paraformaldehyde (PFA) for 2 h at room temperature. Samples were then washed with 1X phosphate-buffered saline (PBS). Embryos were permeabilized in a solution containing 1% Triton X-100, 1% DMSO, and 1% bovine serum albumin (BSA) in PBS for 24 h at 4 °C with gentle agitation. Blocking was performed for an additional 24 h at 4 °C using 10% goat or donkey serum in the same permeabilization solution. Primary antibody incubation was carried out for 4 days at 4 °C in blocking solution using the following antibodies and dilutions: anti–Tenascin-C (#AB19013) Merck (rabbit polyclonal, AB_2256033, 1:200); anti–MF-20 hybridoma bank (mouse monoclonal, AB_2147781, 1:100); anti–Par3 (H-103) Santa Cruz Biotechnology (rabbit polyclonal, 1:100); anti–E-cadherin clone 5D3 hybridoma bank (mouse monoclonal, AB_528116, 1:100); anti–Isl1 clone 39.4D5 hybridoma bank (mouse monoclonal, AB_2314683, 1:200), Collagen 1 clone SP1.D8 hybridoma bank (mouse monoclonal, AB_528438,1:100) and β1-integrin clone 8C8 hybridoma bank (mouse monoclonal, AB_528309, 1:100). After three consecutive washes with 1% Triton X-100 (20 min each), embryos were incubated for 48 h at 4 °C with fluorescent secondary antibodies Donkey anti-Mouse IgG (H+L), Alexa Fluor 555 (#31570) Invitrogen (AB_2536180, 1:300) and Donkey anti-Rabbit IgG (H+L), Alexa Fluor 647 (#31573) Invitrogen (AB_2536183, 1:300), and protected from light. Nuclear staining was performed using Hoechst dye (1:1000). Final washes with 1% Triton X-100 were performed for 24 h.

Embryos were embedded in 1% agarose in distilled water and cut into small cubes for handling. Optical clearing was achieved using a graded methanol series (25%, 50%, 75%, and 100%; 1 h each), followed by incubation in 50% and then 100% benzyl alcohol/benzyl benzoate (BABB) for 20 min each. Cleared samples were imaged using confocal microscopy, Leica DMI8, or light-sheet microscopy, ZEISS Lightsheet 7.

### Protein extraction

Embryos at stages 35 and 42 were lysed in 50 μL of RIPA buffer supplemented with protease inhibitors (Sigma-Aldrich #P8340) for total protein extraction. Embryos were mechanically homogenized using a sterile plastic pestle and centrifuged at 4 °C for 20 min at maximum speed. The supernatant was carefully collected, avoiding both the pellet and the upper lipid layer. Protein concentration was determined using the Pierce™ BCA Protein Assay Micro Kit (Thermo Scientific, #23225), following the manufacturer’s instructions. Measurements were performed in triplicate in 96-well plates.

### Western blot analysis

Proteins were separated by SDS–PAGE using 7.5% polyacrylamide gels and transferred onto nitrocellulose membranes. Membranes were blocked with 5% non-fat milk or BSA in TBS- T and incubated with primary antibodies overnight at 4 °C. After washing, membranes were incubated with HRP-conjugated secondary antibodies for 1 h at room temperature. Protein detection was performed using the SuperSignal™ West Pico PLUS chemiluminescent substrate (Thermo Scientific, #34580), and images were acquired with an iBright Imaging System 1500 (Invitrogen).

### Morpholino microinjection

Antisense morpholinos targeting *Xenopus laevis* fibronectin were designed based on previously published sequences (Davidson et al., 2006) and synthesized by Gene Tools. The morpholino sequences used were: XFN1.MO, 5ʹ- CGCTCTGGAGACTATAAAAGCCAAT-3ʹ; XFN2.MO, 5ʹ- CGCATTTTTCAAACGCTCTGAAGAC-3ʹ.

Embryos at 4-cell stage were injected with 10 or 20 ng of morpholino per cell. A standard control morpholino 5’-CCTCTTACCTCAGTTACAATTTATA-3.’ was used as a negative control. Microinjections were performed in 0.1X MMR supplemented with 5% Ficoll. Two hours after injection, embryos were carefully transferred to 0.1X MMR and grown at 23° C until the desired developmental stage was reached.

### Bioinformatic analysis

Protein sequences of the two *Xenopus laevis* TnC homeologs (Tnc.L, UniProt ID: A0A8J0TG81; Tnc.S, UniProt ID: A0A8J0TQA1) and the *Mus musculus* ortholog (TENA_MOUSE, UniProt ID: Q80YX1) were retrieved from UniProt in FASTA format. Multiple sequence alignment was performed using Clustal Omega with default parameters. Conserved motifs and modular domains were identified using ScanProsite, and the results were compared with available structural annotations in UniProt.

### Image analysis

Image processing and three-dimensional reconstruction were performed using Imaris Viewer. Morphometric measurements, including area and circularity, and digital sectioning of regions of interest, were performed in ImageJ/Fiji. Circularity was calculated as 4 𝜋 × Area / Perimeter ^2^, where a value of 1 corresponds to a perfect circle. Final figures and schematic representations were generated using Adobe Illustrator.

### Statistical analysis

Statistical analyses were performed using GraphPad Prism. An unpaired t-test was applied for comparisons between two groups, while one-way ANOVA was used for multiple comparisons. A p-value < 0.05 was considered statistically significant. Data are presented as mean ± standard deviation of the mean (SD) of at least 3 independent experiments.

## Results

### Second heart field progenitors within the dorsal pericardial wall undergo stage-dependent expansion and cellular stratification in *Xenopus laevis*

The OFT undergoes complex morphogenetic remodeling during early heart development and is frequently affected in congenital heart defects. Because early cardiac development in mammals occurs in utero, making direct observation technically challenging, we compared heart development in *Mus musculus* and *Xenopus laevis*, which undergoes external embryonic development and allows direct access to early stages. To compare cardiac OFT morphology across species, we analyzed the three-dimensional organization of the embryonic heart in mouse embryos at E9.5 and in *Xenopus laevis* at NF35 and NF42, stages at which heart development has been previously described (Kolker et al., 2000; Mohun et al., 2000), using the myocardial marker MF20 (Bader et al., 1982), and 3D reconstruction (Figure 1). MF20 immunostaining showed a well-defined ventricle and OFT in both species, with comparable spatial organization along the proximal–distal axis. In mouse embryos at E9.5, the OFT is clearly subdivided into proximal and distal regions, whereas in *Xenopus* an anatomically equivalent OFT structure was observed at NF42, while this subdivision was not observed at NF35 (Figure 1A-F, movies 1-2). Sagittal sections further confirmed the conserved arrangement of the OFT relative to the ventricle and atrium, as well as their spatial localization adjacent to pharyngeal endodermal and mesodermal tissues (Figure 1G-I, movies 1-2).

**Figure 1.**
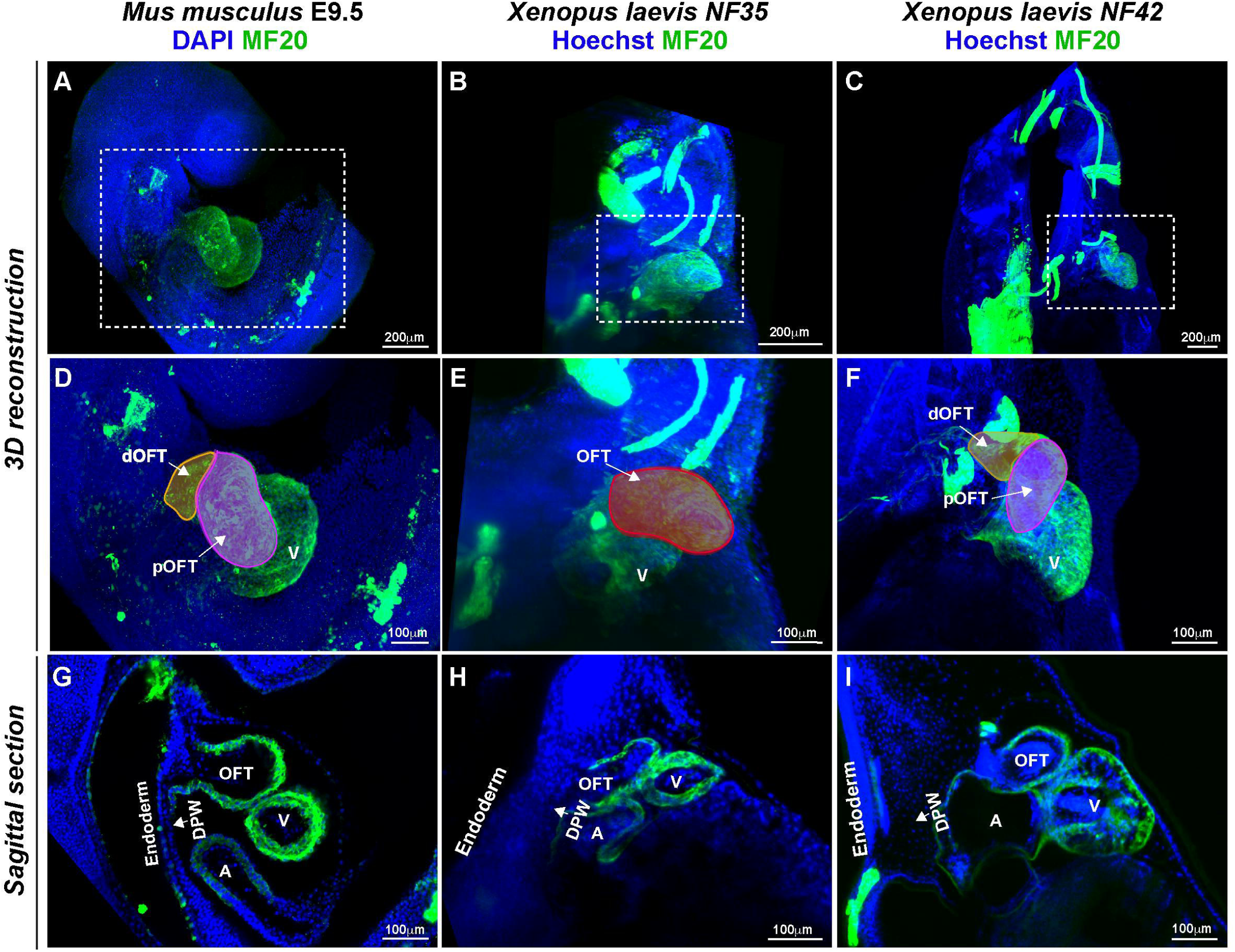
Comparison of the outflow tract elongation during cardiac development in *Mus musculus* and *Xenopus laevis*. **A-I.** 3D reconstructions (**A–F**) and representative sagittal sections (**G–I**) of the embryonic heart in *Mus musculus* at E9.5 and in *Xenopus laevis* at NF35 and NF42 stages. Nuclei were labeled with DAPI or Hoechst (blue), and myocardium was visualized with MF20 (green). In *Mus musculus* E9.5 embryos (**D**), white arrows indicate the proximal (pOFT) and distal (dOFT) regions of the OFT. In *Xenopus laevis*, comparable cardiac structures are observed at NF42 (F). Sagittal sections highlight the relative arrangement of the OFT, ventricle (V), and atrium (A), revealing a conserved tissue organization between species despite differences in developmental timing. The main labeled structures: V, ventricle; A, atrium; OFT, outflow tract (red); dOFT, distal outflow tract (orange); pOFT, proximal outflow tract (magenta); DPW, dorsal pericardial wall; Endoderm, pharyngeal endoderm. (n=4).

Notably, Hoechst nuclear labeling revealed that at NF35 the DPW appeared to comprise predominantly a single cellular layer, resembling the organization observed in mouse embryos (Figure 1G, H, white arrows). In contrast, at NF42, the DPW displayed multiple apparent cell layers, suggesting a more complex tissue architecture at later stages (Figure 1G, white arrows).

In mouse embryos, SHF progenitors reside within the DPW, from which they are progressively incorporated into the elongating OFT (Kelly et al., 2001; Cai et al., 2003; Waldo et al., 2005). Given the stage-dependent differences observed in DPW cellular organization in *Xenopus laevis*, we next sought to further characterize DPW cells at NF35 and NF42, with particular emphasis on determining whether they correspond to SHF progenitor cells. To this end, we performed whole-mount immunostaining to ISL1, a marker of SHF progenitor cells (Cai et al., 2003), and E-cadherin to delineate the adjacent pharyngeal endoderm epithelium (Cortes et al., 2018; Alfano et al., 2019) (Figure 2). At NF35, ISL1+ nuclei were detected both in the endoderm and within a DPW domain predominantly organized as a single cellular layer positioned between the endoderm and the developing heart (Figure 2A-B1, E-E1). In contrast, at NF42 the DPW exhibited increased cellular stratification, with nuclei arranged in multiple layers across the tissue and a noticeable increase in DPW thickness. This change was accompanied by a higher number of ISL1⁺ cells within this expanded region (Figure 2C-D1, F-F1). E-cadherin labeling consistently marked the pharyngeal endoderm at both stages, indicating that the observed increase in cellular layering reflects an expansion of SHF cells within the DPW (Figure 2E-F).

**Figure 2.**
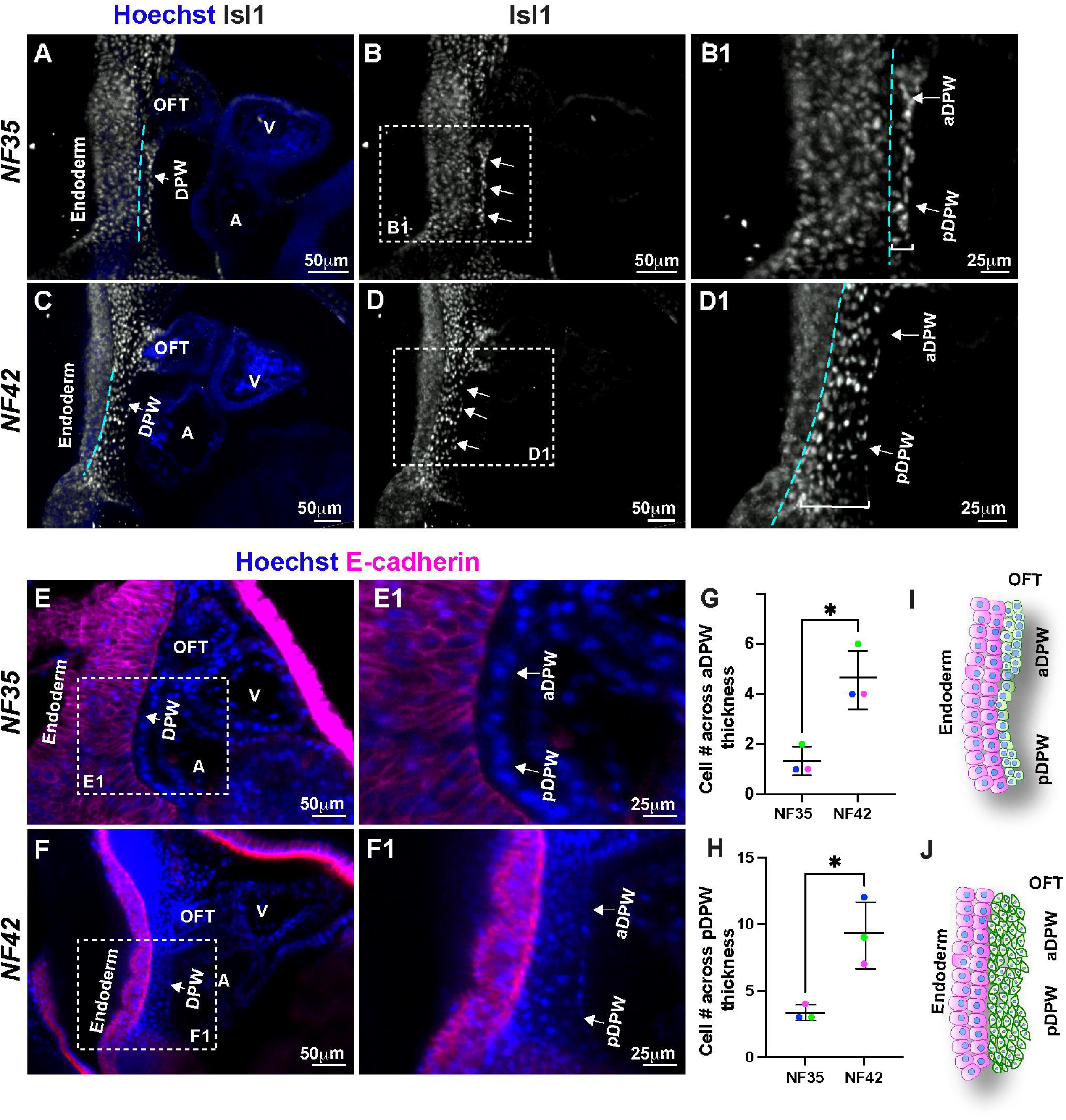
Stage-dependent stratification of SHF-associated cells within the dorsal pericardial wall during *Xenopus laevis* heart development. **A–D1.** NF35 (**A**-**B1**) and NF42 (**C**-**D1**) *Xenopus laevis* embryos were processed by whole-mount immunostaining to detect ISL1 (white) and nuclei (Hoechst, blue). The regions outlined in **B** and **D** are shown at higher magnification in **B1** and **D1,** highlighting the aDPW and pDPW regions. Cyan-dashed lines indicate the boundary between endoderm and DPW. DPW thickness is indicated by the white bracket. **E–F1** NF35 (**E**-**E1**) and NF42 (**F**-**F1**) embryos were processed by whole-mount immunostaining to detect E-cadherin (magenta) and nuclei (Hoechst, blue). The boxed regions in **E** and **F** were magnified in **E1** and **F1**, highlighting the aDPW and pDPW regions. **G**-**F**. Quantification of ISL1-positive nuclei across the thickness of the aDPW and pDPW at NF35 and NF42, (n=3). **I–J**. Schematic representations summarizing the stage-dependent transition of the DPW from a predominantly monolayered organization at NF35 (**I**) to a multilayered structure at NF42 (**J**) while maintaining its spatial relationship with the endoderm and the developing OFT. Main labeled structures: V, ventricle; A, atrium; OFT, outflow tract; dOFT, distal outflow tract; pOFT, proximal outflow tract; aDPW, anterior dorsal pericardial wall ; pDPW, posterior dorsal pericardial wall ; Endoderm, pharyngeal endoderm.

To further characterize changes in DPW cellular organization across stages and to facilitate comparison with mammalian development, we quantified the number of ISL1⁺ nuclei across the thickness of the dorsal pericardial wall at NF35 and NF42. Because the anterior and posterior domains of the DPW contribute to distinct regions of the developing OFT (Bertrand et al., 2011; Francou et al., 2013), these domains were analyzed separately. We observed a significant increase in the number of ISL1⁺ nuclei across the thickness of the anterior DPW (aDPW) at NF42 compared with NF35 (Figure 2G). A similar increase was detected in the posterior DPW (pDPW), where NF42 embryos also exhibited a marked rise in the number of nuclei relative to NF35 (Figure 2H). These observations suggest a progressive expansion and redistribution of SHF progenitors within the DPW between NF35 and NF42.

Together, these results suggest that in *Xenopus laevis*, SHF cells localized in the DPW undergo a stage-dependent transition from a predominantly monolayered organization at NF35 to a multilayered structure at NF42, while preserving their spatial relationship with the endoderm and the developing OFT (Figure 2I–J).

### SHF-associated cells in the DPW undergo stage-dependent remodeling of epithelial organization and cell shape during heart development

Previous studies in mouse embryos have established that SHF progenitors localized in the DPW exhibit epithelial characteristics that are critical for heart tube elongation, including cell polarity, adhesion, and tissue-level mechanical properties (Francou et al., 2014, 2017; Cortes et al., 2018). Since we observed a stage-dependent increase in cellular stratification of SHF-associated cells in the *Xenopus laevis* DPW (Figure 2), we next asked whether this transition was accompanied by changes in epithelial organization and cell shape. To address this, we performed whole-mount immunostaining for β1-integrin. Integrins are α/β heterodimeric transmembrane receptors composed of α and β subunits that mediate cell-ECM interactions (Sun et al., 2016). We observed that β1-integrin was enriched at the interface between the endoderm and the DPW (Figure 3A1–B1, yellow arrows), consistent with previous observations in mouse (Arriagada et al., 2025). Additionally, we used β1-integrin to delineate DPW cell boundaries, thereby enabling assessment of tissue organization and cell morphology. At NF35, β1-integrin labeling showed that SHF cells showed an epithelial-like organization (Figure 3A–A1, white arrows). In contrast, at NF42, DPW cells exhibited increased spacing between adjacent cells compared with NF35 (Figure 3B–B1, white arrows).

**Figure 3.**
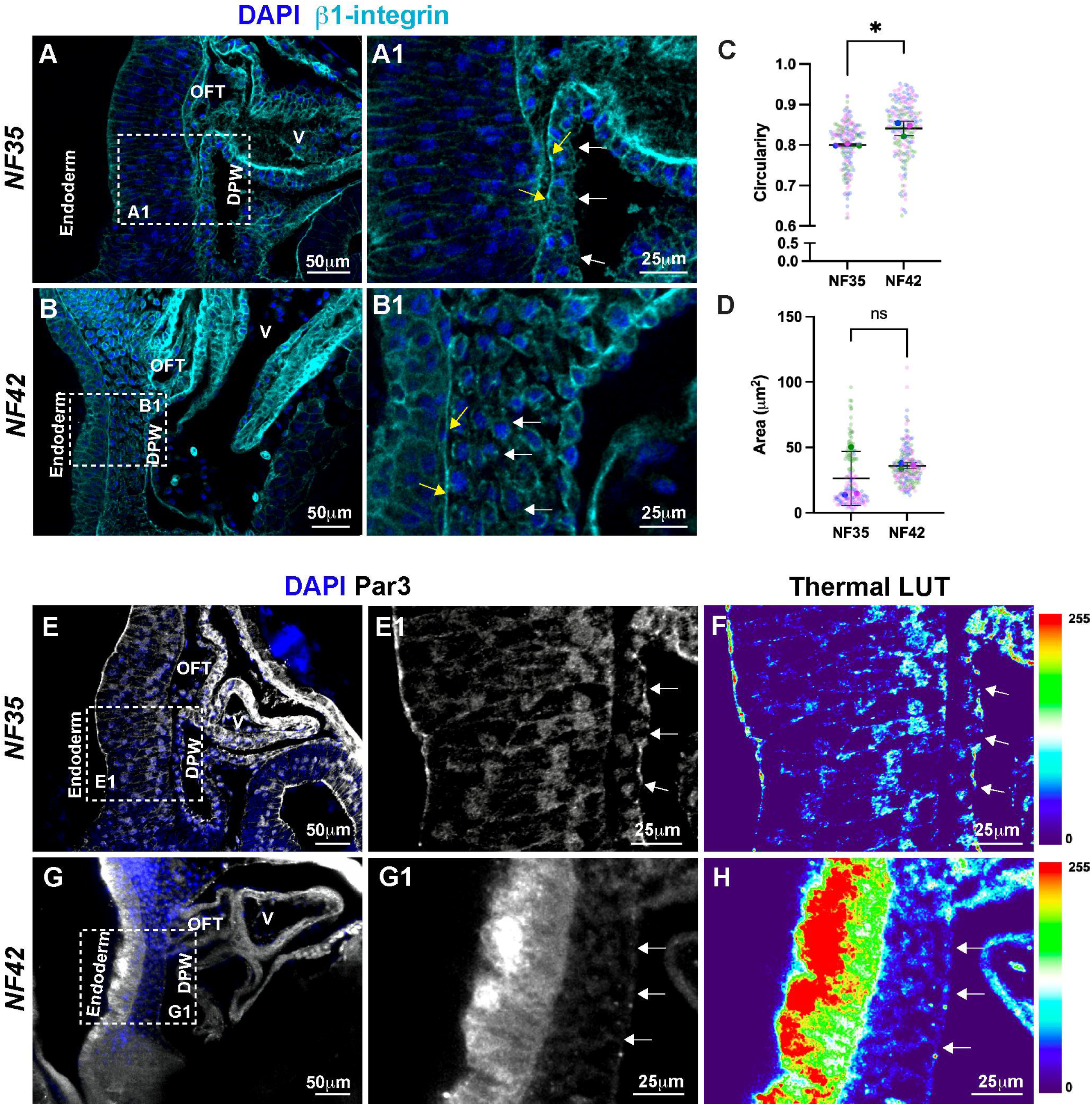
Stage-dependent changes in the DPW epithelial organization, cell shape, and polarity. **A–B**. Whole-mount immunostaining of *Xenopus laevis embryos* at NF35 (A) and NF42 (B) labeled for β1-integrin (cyan) and nuclei (DAPI, blue). Dashed boxes in A and B indicate the DPW area magnified in A1 and B1. β1-integrin prominently delineates the boundaries of DPW cells (white arrows). Yellow arrows indicate integrin-enriched interfaces facing the pharyngeal endoderm. **C-D**. Quantification of DPW cell area and circularity at NF35 compared to NF42 (C). Each dot represents an individual embryo; bars indicate mean ± SD. p < 0.05; ns, not significant. **E–H.** Analysis of epithelial polarity using Par3 immunostaining (white) with DAPI (blue). At NF35 (E, E1) and NF42 (G, G1). F, H. Thermal LUT representations highlight increased Par3 signal intensity.

Because β1-integrin clearly marked cell boundaries at both stages, this staining was used to quantify cell-shape parameters. Elongated cells exhibit circularity values near 0, whereas round cells exhibit circularity close to 1. Cell circularity was significantly higher at NF42 than at NF35 (Figure 3C), indicating a stage-dependent shift toward a more circular cell shape during DPW remodeling. However, quantitative analysis showed no significant differences in cell area between NF35 and NF42 DPW cells (Figure 3D).

To further examine epithelial organization and polarity in DPW cells, we analyzed the localization of Par3, a key component of the apical polarity complex (Thompson, 2022). At NF35, Par3 exhibited continuous apical membrane enrichment in DPW cells, consistent with a well-organized epithelial architecture (Figure 3E–F, white arrows). In contrast, at NF42, Par3 localization appeared as a more discontinuous enrichment of Par3 (Figure 3G–H, white arrows).

Together, these findings demonstrate that SHF progenitors cells in the *Xenopus laevis* DPW undergo stage-dependent epithelial remodeling characterized by changes in cell shape, spacing, and polarity

### ECM remodeling underlies DPW architectural changes during heart tube elongation in *Xenopus laevis*

Given that epithelial remodeling of the DPW was accompanied by changes in cell shape, cell–cell spacing, and apico-basal polarity, we next asked whether this cellular reorganization was associated with concomitant changes in the ECM composition surrounding SHF-associated cells. The epithelial architecture and polarity are tightly regulated by cell–ECM interactions, and ECM components provide both structural support and mechanical cues during heart tube elongation (Rozario and DeSimone, 2010; Lockhart et al., 2011; Arriagada et al., 2025). We therefore asked whether DPW maturation is accompanied by stage-dependent modifications in ECM composition and distribution. To address this, we analyzed the temporal expression and spatial distribution of key ECM components implicated in SHF morphogenesis and OFT elongation, including Fn1, TnC. ColI and Laminin (Lam) (Löhler et al., 1984; Yarnitzky and Volk, 1995; Rahkonen et al., 2004; Astrof et al., 2007; Derrick et al., 2021; Arriagada et al., 2025).

Western blot analysis revealed a progressive, stage-dependent increase in Fn1 protein levels from NF14 to NF42 (Figure 4A–B). In contrast, TnC and Col I were barely detected at early stages (NF14–NF30) and showed a marked upregulation at NF35 and NF42 (Figure 4A, C– D), coinciding with developmental stages at which DPW epithelial organization and cell morphology undergo significant changes. These results suggest that DPW maturation is accompanied by a coordinated temporal remodeling of the ECM.

**Figure 4.**
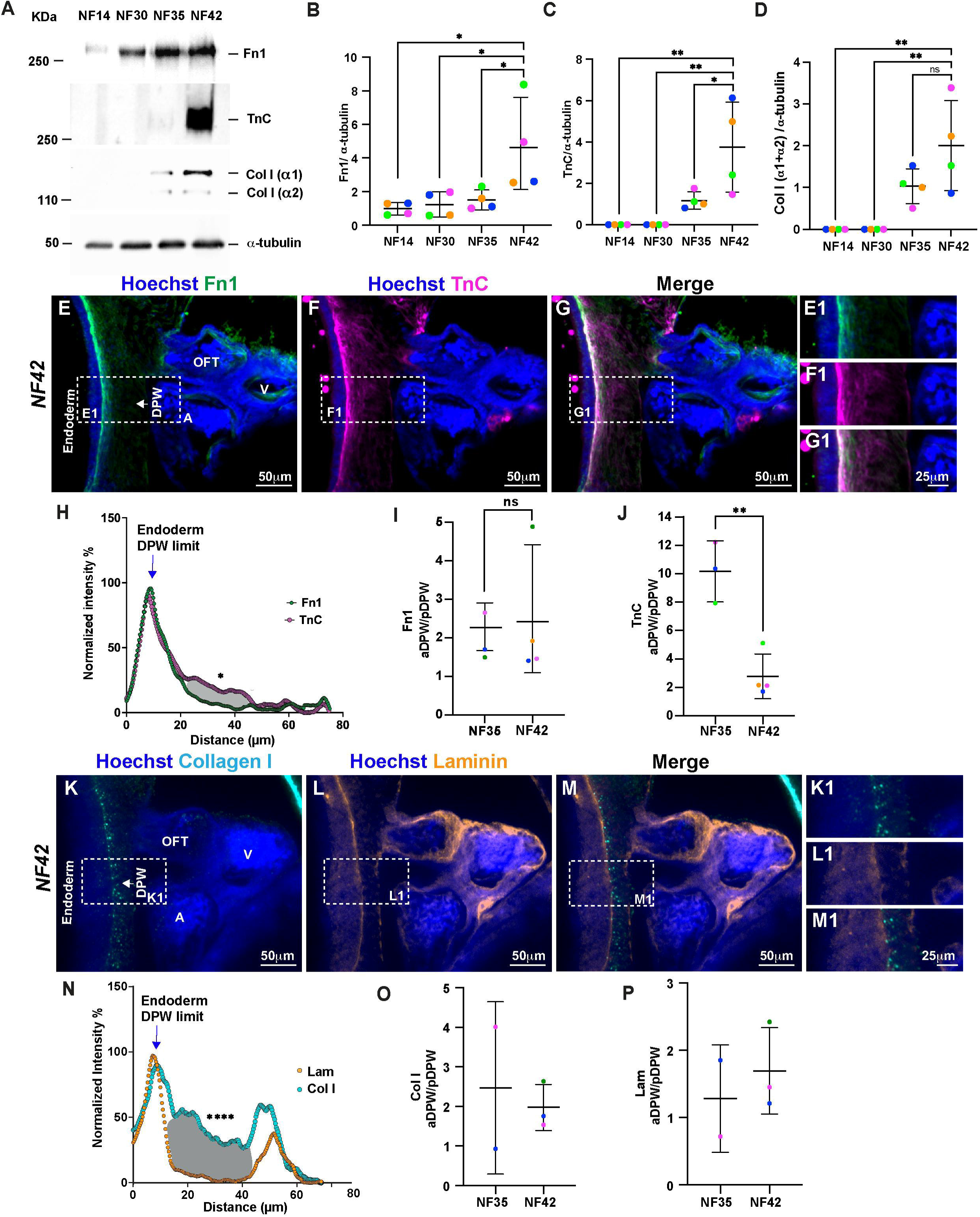
Stage-dependent remodeling of ECM composition and spatial distribution in the dorsal pericardial wall during Xenopus laevis heart development. **A.** Representative Western blot analysis of Fibronectin 1 (Fn1), Tenascin-C (TnC), and Collagen I (Col I α1 and α2 chains) protein levels in whole embryos at developmental stages NF14, NF30, NF35, and NF42. α-tubulin was used as a loading control. **B–D.** Densitometric quantification of Fn1 (B), TnC (C), and Col I (α1+α2) (D) protein levels normalized to α-tubulin. Each dot represents an individual biological replicate. **E–G.** Whole-mount immunostaining of NF42 embryos showing Fn1 (green), TnC (magenta), and merged signal, with nuclei labeled with Hoechst (blue). Dashed boxes indicate the dorsal pericardial wall (DPW) region adjacent to the pharyngeal endoderm and outflow tract (OFT). **E1–G1.** Higher-magnification views of boxed regions illustrating Fn1 enrichment along tissue boundaries and TnC localization within the DPW. **H.** Line-scan intensity profiles across the DPW at NF42 comparing normalized Fn1 and TnC fluorescence signals, illustrating distinct spatial distributions of these ECM components **I–J.** Quantification of fluorescence intensity ratios between anterior and posterior DPW (aDPW/pDPW) for Fn1 (H) and TnC (I) at NF35 and NF42. **K–M.** Whole-mount immunostaining of NF42 embryos showing Collagen I (cyan), Laminin (orange), and merged signal, with nuclei labeled with Hoechst (blue). Dashed boxes indicate the DPW region. **K1–M1.** Higher-magnification views highlighting robust accumulation of Collagen I surrounding the DPW and a more restricted Laminin distribution associated with basement membrane–like structures. **N.** Representative line-scan intensity profiles across the DPW at NF42 comparing Laminin and Collagen I distributions. **O–P.** Quantification of aDPW/pDPW fluorescence intensity ratios for Collagen I (O) and Laminin (P) at NF35 and NF42. Main labeled structures: V, ventricle; A, atrium; OFT, DPW, dorsal pericardial wall ; Endoderm, pharyngeal endoderm.

To assess whether changes in the spatial organization of ECM proteins occur within the DPW, we examined their distribution using whole-mount immunostaining. At NF35, Fn1 and TnC exhibited distinct yet partially overlapping distributions along the DPW–endoderm interface (Supplementary Figure S1). Notably, Fn1 was continuously distributed along the DPW, in contrast to mouse embryos, where Fn1 is predominantly localized at the DPW–endoderm interface (Arriagada et al., 2025). In contrast, TnC displayed a more restricted pattern, largely confined to the endoderm and adjacent DPW region, similar to that of the mouse.

At NF42, Fn1 remained detectable along tissue boundaries at the DPW–endoderm interface (Figure 4E–E1); however, its intercellular localization within the DPW was markedly reduced in comparison to NF35 (Figure S1, 4E-E1). In contrast, TnC exhibited a pronounced redistribution at NF42, with increased accumulation within the endoderm and DPW, and enhanced localization between DPW cells (Figure S1, 4F-F1). When we analyzed the distribution of Fn1 and TnC, we observed that they are co-distributed at the endoderm–DPW boundary but display distinct localization patterns within the DPW, with TnC being more prominently enriched between SHF cells than Fn1 (Figure 4H). Quantitative analysis of the fluorescence intensity ratio between the anterior and posterior DPW (aDPW/pDPW) reveals that the Fn1 ratio remained largely unchanged (Figure 4I), while there was a significant reduction in the TnC ratio at NF42 compared with NF35 (Figure 4J), indicating a region-specific redistribution.

We next examined fibrillar and basement membrane-associated ECM components at NF42. ColI displayed robust accumulation surrounding the DPW and adjacent mesenchyme (Figure 4K-K1, N), whereas Lam showed a more confined distribution, largely restricted to basement membrane-like structures at the DPW–endoderm boundary and the apical region of the DPW (Figure 4M-M1, N), with no changes in the aDPW/pDPW distribution (Figure 4O-P).

Together, these data reveal a stage-dependent reorganization of ECM composition within the DPW, characterized by a reduced Fn1 and an increased TnC intercellular localization at NF42, accompanied by enhanced fibrillar ECM deposition. This redistribution likely contributes to DPW remodeling and the associated changes in tissue architecture during OFT tract development.

Because the localization pattern of TnC in *Xenopus laevis* differs from that described in mouse, we performed an *in silico* comparative analysis of sequence conservation and domain organization across vertebrates (Supplementary Figure S2A). Sequence alignment and domain prediction confirmed that both *Xenopus laevis* homeologs retain the conserved N-terminal assembly domain, EGF-like repeats, constitutive FNIII domains, and the C-terminal fibrinogen-like globe (Supplementary Figure S2B). Differences relative to mouse were restricted to the central FNIII domains (Supplementary Figure S2B, orange domains), which are encoded by alternatively spliced exons and are known to vary across vertebrate species. Despite this variability, the core modular organization of TnC remains conserved. Thus, the differences between the mouse and *Xenopus* TnC sequence are within the alternatively spliced cassette rather than loss of essential structural domains.

### Fn1 depletion impairs outflow tract elongation and ventricular morphogenesis

Fn1 is a key ECM component that supports embryonic tissue remodeling by regulating cell– matrix adhesion, collective cell behavior, and force transmission through integrin-dependent mechanisms (Hynes, 1986; Pankov and Yamada, 2002). Genetic and functional studies across vertebrate models have demonstrated that Fn1 is essential for processes requiring extensive tissue remodeling, including gastrulation, somitogenesis, and organogenesis (George et al., 1993; Davidson et al., 2006; Astrof et al., 2007a). Additionally, in mouse models, Fn1 has been implicated in the deployment of cardiac progenitors, myocardial differentiation, and OFT elongation (Arriagada et al., 2025). To determine whether Fn1 is required for cardiac morphogenesis in a manner conserved across vertebrates, we examined the effects of Fn1 depletion on OFT and ventricular development in *Xenopus laevis*

For knocking down Fn1, different concentrations of a translation-blocking Fn1 morpholino (Fn1-MO) were injected at the 2-4 cell stage, and Fn1 protein levels were assessed at 48- and 72-hours post-fertilization (hpf), corresponding to NF35 and NF42, respectively. As expected, Fn1 protein levels were significantly reduced in embryos injected with either 10 ng or 20 ng of Fn1 morpholino (Fn1-MO) compared with control morpholino (Co-MO) embryos at both 48 and 72 hpt (Figure 5A–C). This reduction confirms an effective decrease in Fn1 expression throughout the period of cardiac morphogenesis.

**Figure 5.**
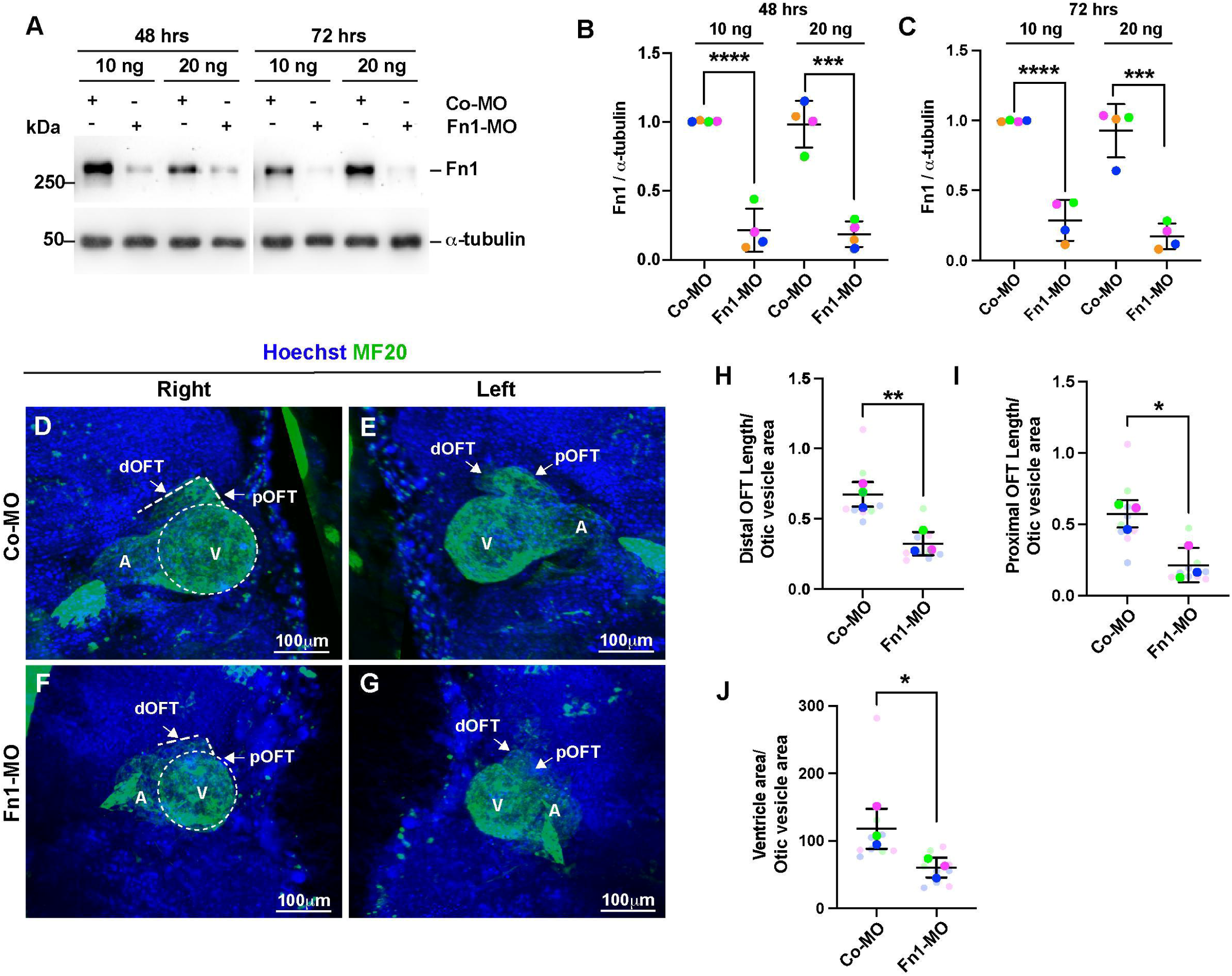
Fn1 depletion impairs outflow tract elongation and ventricular morphogenesis in Xenopus laevis. **A.** Western blot analysis of Fn1 protein levels in embryos injected with control morpholino (Co-MO) or Fn1 morpholino (Fn1-MO) at 48 and 72 hours post-fertilization (hpf). Embryos were injected with 10 ng or 20 ng of morpholino as indicated. α-tubulin was used as a loading control. **B–C.** Densitometric quantification of Fn1 protein levels normalized to α-tubulin. Each dot represents an individual biological replicate. **D–G.** Whole-mount immunostaining for MF20 (green) with Hoechst nuclear labeling (blue) showing cardiac morphology in control (**D–E**) and Fn1-depleted (**F–G**) embryos at NF42. Representative right and left views are shown. Dashed outlines indicate the ventricle (V), and arrows mark the distal (dOFT) and proximal (pOFT) regions of the outflow tract. A, atrium. **H–I.** Quantification of distal (**H**) and proximal (**I**) OFT length normalized to the otic vesicle area. **J.** Quantification of ventricular area normalized to the otic vesicle area. Data are presented as mean ± SD. Statistical significance was determined using t-test; *p* < 0.05, **p**< 0.01, ***p*** < 0.001, **p** < 0.0001.

We next assessed cardiac morphology by whole-mount MF20 immunostaining. Control embryos exhibited a well-extended OFT with clearly distinguishable pOFT and dOFT domains and proper alignment relative to the ventricle and atrium (Figure 5D–E). In contrast, Fn1-depleted embryos displayed a markedly shortened OFT, characterized by reduced extension of both proximal and distal regions and a more compact myocardial configuration (Figure 5F–G).

Quantitative analyses confirmed a significant reduction in distal and proximal OFT length in Fn1-MO embryos compared to controls (Figure 5H–I). In addition, Fn1 depletion resulted in a significant decrease in ventricular area (Figure 5J), indicating that Fn1 is required for proper OFT elongation and ventricular morphogenesis. These defects suggest a conserved function of Fn1 during cardiac development.

### Fn1 depletion alters ECM composition during cardiac development

Given the established role of Fn1 in organizing the ECM (Sottile and Hocking, 2002), we next asked whether Fn1 depletion affected the abundance and organization of other key ECM components during cardiac morphogenesis.

To address Fn1 effect on ECM protein abundance, we examined the effects of Fn1 knockdown on TnC and ColI protein levels at 48 and 72 hpf. Western blot analysis revealed that Fn1 depletion altered ECM protein levels (Figure 6A). TnC protein levels were not significantly changed in Fn1-MO embryos injected with 10 ng; however, a significant reduction in TnC levels was observed at the higher Fn1-MO dose (20 ng) at both time points (Figure 6B). In contrast, ColI levels were affected at both MO concentrations and at both time points evaluated. Quantification of the ColI α1 chain revealed a significant reduction in Fn1-MO embryos at both 48 and 72 hpf, even at the lower morpholino dose (Figure 6C). Similarly, the ColI α2 chain was significantly decreased at 48 hpf in Fn1-depleted embryos; although no differences were observed at 72 hpf (Figure 6D). Together, these data suggest that Fn1 is required for the proper expression of ECM proteins, with ColI appearing more sensitive to Fn1 depletion than TnC.

**Figure 6.**
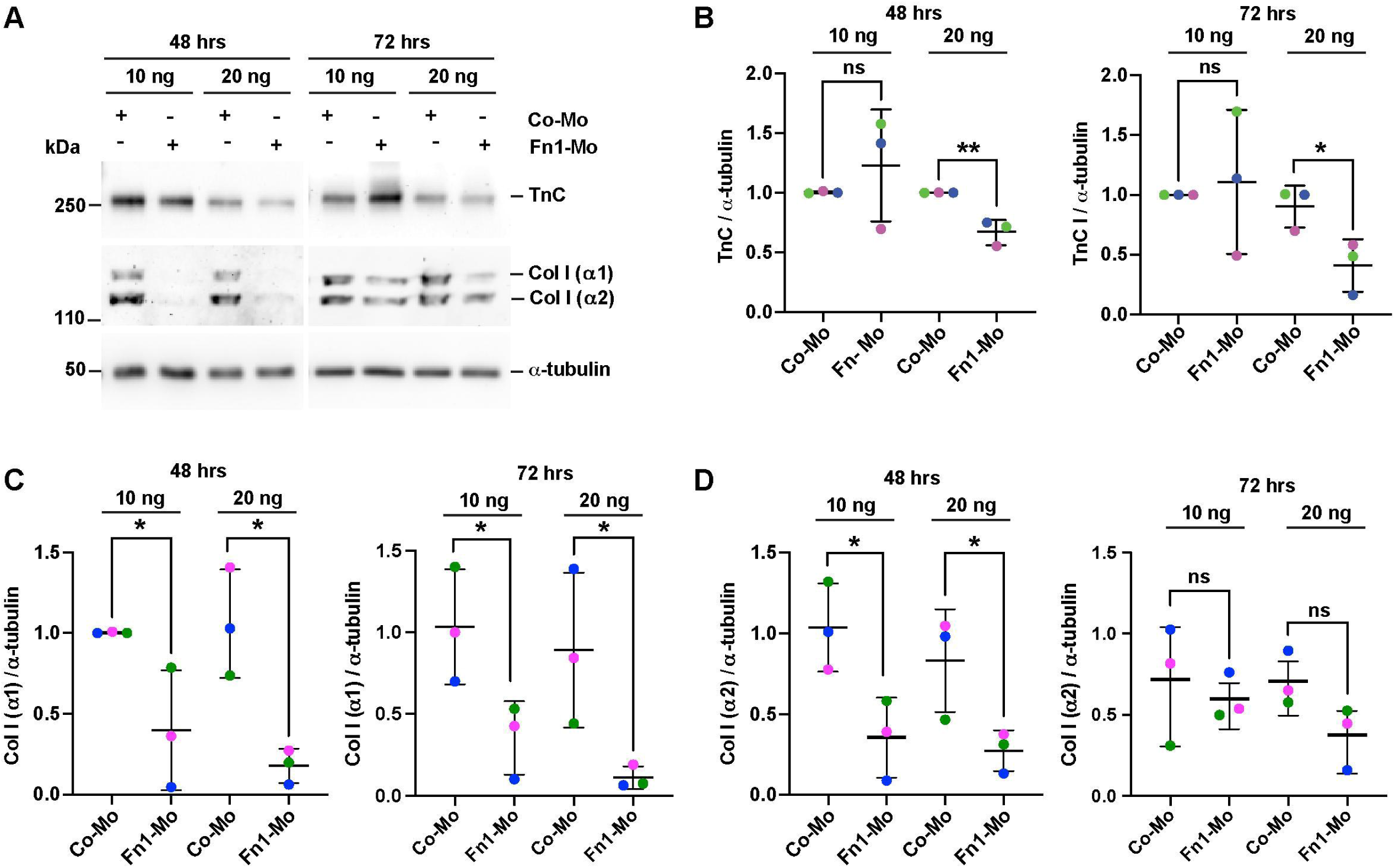
Fn1 depletion alters ECM composition. **A.** Representative Western blot analysis of TnC and Collagen I (Col I α1 and α2 chains) protein levels in embryos injected with control morpholino (Co-MO) or Fn1 morpholino (Fn1-MO) at 48 and 72 hours post-fertilization (hpf). Embryos were injected with 10 ng or 20 ng of morpholino as indicated. α-tubulin was used as a loading control. **C.** Densitometric quantification of TnC protein levels at 48 hpf (left) and 72 hpf (right), expressed in arbitrary units (a.u.) and normalized to α-tubulin. **D.** Densitometric quantification of Collagen I α1 chain levels normalized to α-tubulin at 48 hpf (left) and 72 hpf (right). **(E)** Densitometric quantification of Collagen I α2 chain levels normalized to α-tubulin at 48 hpf (left) and 72 hpf (right) Each dot represents an individual biological replicate. Data are presented as mean ± SD. Statistical significance was determined using t-tests; *p* < 0.05, **p** < 0.01; ns, not significant

To assess whether Fn1 depletion alters the spatial organization of TnC within the DPW, we performed whole-mount immunostaining for Fn1 and TnC at NF42, as at this stage the OFT is fully elongated, and we previously observed a higher abundance of ECM components within the DPW. In control embryos, Fn1 formed a continuous fibrillar network along the DPW–endoderm interface and around the outflow tract, while TnC displayed a homogeneous signal within the DPW region (Figure 7A–C, C1, arrows). In contrast, Fn1-depleted embryos showed the expected disruption of the Fn1 fibrillar network together with a marked reduction and discontinuity of TnC signal within the DPW (Figure 7D–F, F1, arrows).

**Figure 7.**
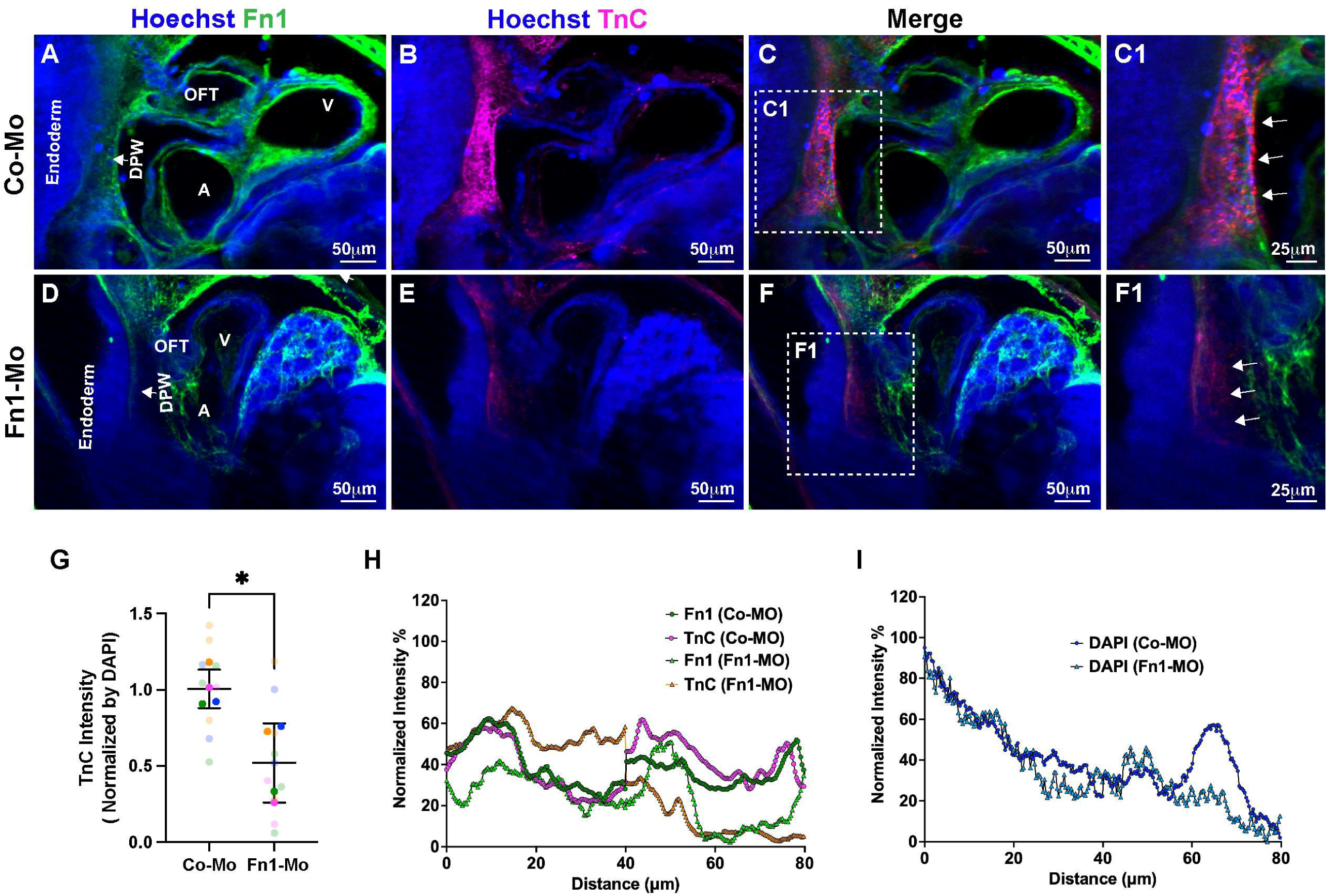
Fn1 depletion decreases TnC expression in the DPW. **A–F1.** Whole-mount immunofluorescence of NF42 embryos injected with control morpholino (Co-MO) (A-C1) and Fn1 morpholino (Fn1-MO), showing Fn1(green), TnC (magenta), and nuclei labeled with Hoechst (blue). Fn1 and TnC are detected in the dorsal pericardial wall (DPW), adjacent to the pharyngeal endoderm and the outflow tract (OFT). **C1, F1.** Higher-magnification view of the boxed region in (C) highlighting intercellular localization of TnC within the DPW (arrows). **G.** Quantification of TnC fluorescence intensity within the DPW, normalized to DAPI signal. **H.** Representative line-scan intensity profiles across the DPW illustrating the spatial distribution of Fn1 and TnC in control and Fn1-depleted embryos. **I.** Representative line-scan intensity profiles across the DPW illustrating the spatial distribution of DAPI in control and Fn1-depleted embryos Each dot represents an individual biological replicate. Data are presented as mean ± SD. Statistical significance was determined using t-test, < 0.05, p < 0.01; ns, not significant

Quantitative analysis confirmed a significant reduction in TnC intensity in Fn1-depleted embryos compared with controls (Figure 7G). Line-scan intensity profiles across the DPW revealed distinct effects on ECM component distribution following Fn1 depletion. Fn1- depleted embryos showed a marked reduction in Fn1 signal across the entire analyzed region, indicating a global decrease in Fn1 levels. Notably, TnC exhibited a clear redistribution, with altered peak patterns and a loss of the coordinated spatial profile observed under control conditions (Figure 7H). To assess whether these changes were associated with alterations in tissue architecture, we analyzed the distribution of DAPI signal along the same axis. Line-scan analysis revealed a reduced spatial extent of DAPI staining in Fn1-depleted embryos compared to controls, indicating a decrease in DPW width (Figure 7I). Together, these results suggest that Fn1 depletion not only disrupts ECM organization but also affects tissue architecture, leading to a narrowing of the DPW. Together, these data indicate that Fn1 is required to maintain both normal levels and spatial organization of TnC within the SHF-associated DPW.

### Fn1 depletion reduces DPW area without affecting SHF progenitor number or individual cell morphology

Given that Fn1 is known not only to organize ECM architecture but also to play a key role in tissue remodeling (Rozario and DeSimone, 2010), we next examined whether Fn1 depletion affects the organization and cellular properties of the SHF progenitor cells within DPW at NF42.

To assess SHF progenitor identity and abundance, we performed whole-mount immunostaining for ISL1 in control and Fn1 MO embryos (Figure 8A–B1). ISL1⁺ nuclei were detected in the DPW in both conditions and remained spatially associated with the outflow tract and surrounding tissues. Quantification of ISL1⁺ cells was performed within a region of interest (ROI) of a defined area, allowing comparison of cell density between conditions. This analysis revealed no significant difference in the number of ISL1⁺ cells within the ROI between control and Fn1-depleted embryos (Figure 8C), indicating that Fn1 depletion does not affect the disposition of SHF progenitors at this stage.

**Figure 8.**
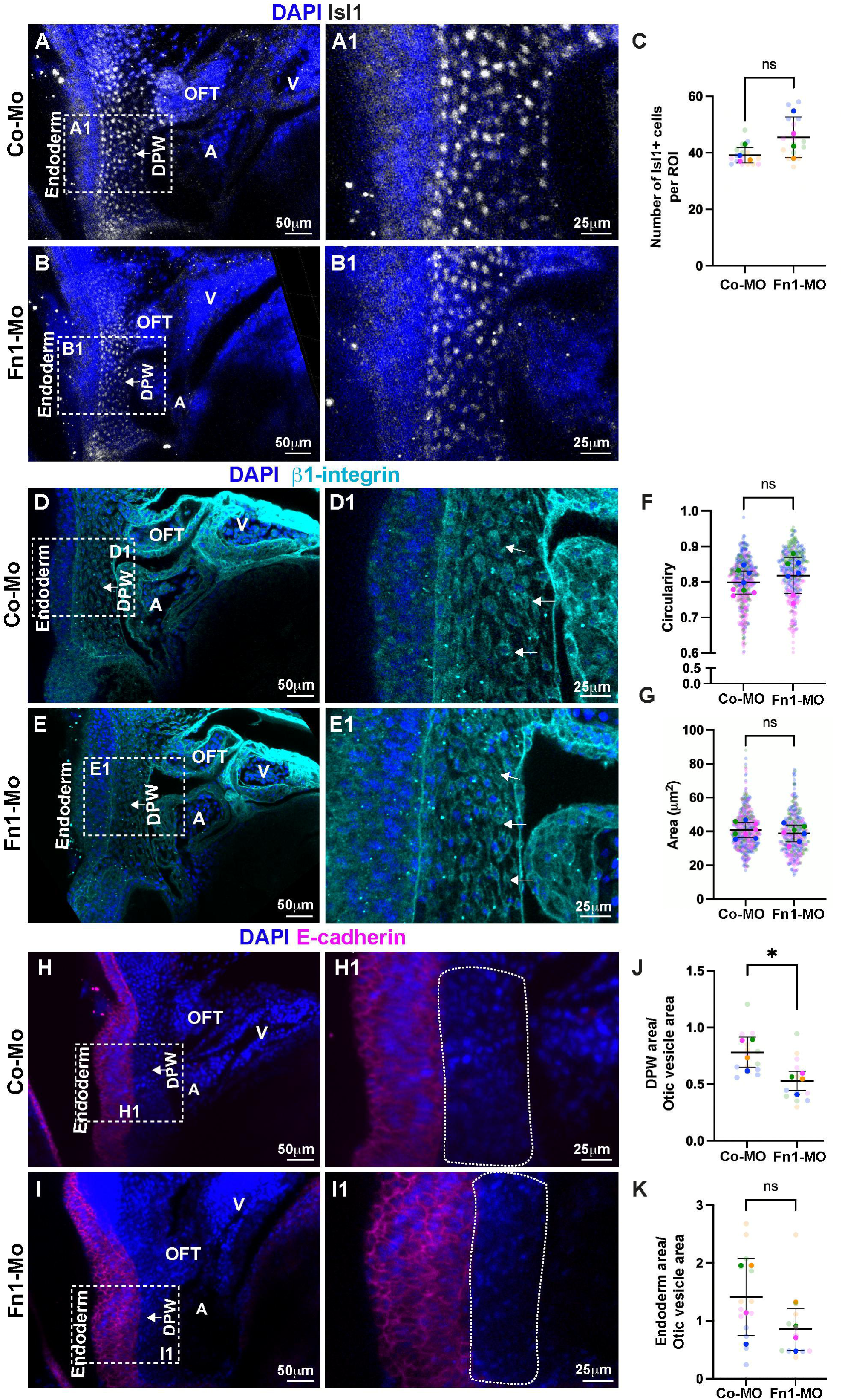
Fn1 depletion reduces DPW tissue area without affecting SHF progenitor number or individual cell morphology. **A–B.** Whole-mount immunofluorescence of NF42 embryos injected with control morpholino (Co-MO) (**A**) or Fn1 morpholino (Fn1-MO) (**B**) stained for ISL1 (white) and nuclei labeled with DAPI (blue). Dashed boxes indicate the dorsal pericardial wall (DPW) region located between the pharyngeal endoderm and the developing outflow tract (OFT). **A1–B1.** Higher-magnification views of the boxed regions showing ISL1⁺ nuclei within the DPW. **C.** Quantification of the number of ISL1⁺ nuclei within the DPW. **D–E.** Whole-mount immunofluorescence of NF42 embryos stained for β1-integrin (cyan) and DAPI (blue) in control (D) and Fn1-depleted (E) embryos. Arrows indicate β1-integrin labeling at DPW cell boundaries. **D1–E1.** Higher-magnification views of the boxed regions. **F–G.** Quantification of DPW cell circularity (F) and cell area (J) based on β1-integrin- defined cell boundaries, showing no significant differences between control and Fn1- depleted embryos. **H–I.** Whole-mount immunofluorescence of NF42 embryos stained for E-cadherin (magenta) and DAPI (blue) to delineate the pharyngeal endoderm and define DPW boundaries in control (H) and Fn1-depleted (I) embryos. **H1–I1.** Higher-magnification views of the boxed regions illustrating DPW morphology. **J- K.** Quantification of DPW area (J) and endodermal area (K), normalized to the otic vesicle area. Data are presented as mean ± SD. Statistical significance was determined using t-tests; *p* < 0.05; ns, not significant.

To determine if there was a change in individual cell morphology, we analyzed DPW cells using β1-integrin immunostaining to delineate cell boundaries (Figure 8D–E1). At NF42, β1- integrin continued to mark DPW cell membranes in both control and Fn1-depleted embryos. Quantitative measurements revealed no significant differences in cell circularity (Figure 8F) or cell area (Figure 8G) between conditions, indicating that Fn1 loss does not appreciably alter individual DPW cell shape or size.

Qualitative observations suggested an apparent decrease in the overall number of ISL1⁺ cells in Fn1-depleted embryos. To determine whether this reflected changes in tissue organization, we analyzed DPW architecture using E-cadherin immunostaining to delineate the adjacent pharyngeal endoderm and define DPW boundaries (Figure 8H–I1). Quantitative analysis confirmed a significant decrease in DPW area in Fn1 MO embryos compared with controls (Figure 8J), whereas the endodermal area remained unchanged (Figure 8K). DPW measurements were normalized to the otic vesicle area to account for potential differences in embryo size. Because ISL1⁺ progenitors are located within the DPW, the reduction in DPW area likely reflects a contraction of the ISL1⁺ progenitor domain rather than differences in overall embryo size. Together, these results indicate that Fn1 depletion reduces the size of the SHF progenitor territory by affecting DPW tissue architecture without altering the local density of ISL1⁺ cells.

Together, these data demonstrate that Fn1 depletion does not impair SHF progenitor identity or individual cell geometry but instead reduces DPW tissue size and the total number of ISL+ cells, accompanied by the concurrent loss of key ECM components. These findings support the idea that Fn1 is required to maintain proper higher-order DPW architecture during late stages of cardiac outflow tract development.

## Discussion

### Stage-dependent ECM remodeling accompanies DPW maturation in Xenopus

The present work describes, for the first time, the cellular organization of the DPW in amphibians. Our data reveal a clear stage-dependent transition in the organization of SHF cells within the DPW between NF35 and NF42. At NF35, the DPW displays a predominantly monolayered organization with relatively few ISL1⁺ cells. By NF42, however, the tissue becomes multilayered and exhibits a marked increase in the number of ISL1⁺ nuclei distributed across its thickness (Figures 1-2), indicating progressive expansion and architectural remodeling of the SHF progenitor cells.

In mammalian embryos, SHF progenitors are typically arranged as a cohesive epithelial layer within the DPW and are progressively incorporated into the elongating OFT rather than forming a multilayered tissue (Francou et al., 2017). Interestingly, our results suggest that these *Xenopus* stages capture distinct aspects of the E9.5 heart. While the degree of OFT elongation observed at NF42 more closely resembles that of mouse embryos at this stage, the overall DPW tissue morphology and spatial distribution of SHF cells more closely resemble the organization observed at NF35 (Kelly et al., 2014).

This architectural transition occurs in parallel with dynamic changes in ECM composition and distribution. Fn1 protein levels progressively increase from NF14 to NF42; however, its spatial organization changes over time. At NF35, Fn1 is detected between DPW cells, whereas at NF42 its intercellular localization is reduced. In contrast, TnC shows enhanced intercellular accumulation at NF42, accompanied by increased ColI deposition (Figure 3). These coordinated changes in ECM distribution indicate that DPW remodeling involves reorganization of matrix architecture rather than a simple quantitative increase in ECM abundance.

Importantly, this organization differs from that described in mouse embryos (Arriagada et al., 2025). In the anterior DPW of the mouse, Fn1 is enriched in the mesoderm and supports epithelial cohesion and mechanotransduction. However, ECM components are primarily localized at basal or tissue-boundary interfaces, particularly at the DPW–endoderm boundary, without forming prominent intercellular accumulations. SHF cells, therefore, maintain a tightly packed epithelial arrangement. In contrast, our findings in *Xenopus* reveal substantial intercellular ECM redistribution at later stages, suggesting species-specific differences in matrix topology that may influence epithelial packing and the mechanical environment.

Additionally, TnC is known to directly bind to the Fn-1 type III repeats, interfering with co-receptor binding (e.g., Syndecan-4) and preventing the maturation of focal adhesions, thereby reducing cytoskeletal tension (Midwood and Schwarzbauer, 2002). As a consequence, TnC can reduce Fn1-mediated adhesion and generate microenvironments with lower adhesive strength (Radwanska et al., 2017). This transition suggests the possibility of a localized Epithelial-to-Mesenchymal Transition (EMT)-like process, driven by TnC-mediated modulation of the Fn1 scaffold, a prerequisite for progenitor deployment into the elongating OFT. The increase in TnC between SHF cells may therefore influence progenitor cells morphology and migratory behavior by altering cell shape, promoting a more rounded morphology, and reducing epithelial-like organization.

Mechanistically, integrin–Fn1 interactions are key regulators of cell polarity, cytoskeletal organization, and tissue mechanics. Integrin-based adhesions are mechanosensitive structures that couple ECM composition to actomyosin tension and to intracellular signaling pathways such as RhoA and YAP/TAZ (Dupont et al., 2011). In the mouse SHF, mesodermal Fn1 maintains epithelial organization and counteracts TnC-mediated anti-adhesive effects, functioning upstream of mechanotransduction (Arriagada et al., 2025). In *Xenopus*, changes in ECM composition are therefore likely to alter adhesion strength and tissue-level tension within the DPW. Early Fn1 enrichment may provide a pro-adhesive scaffold that promotes epithelial cohesion and controlled progenitor deployment, whereas later ECM remodeling may facilitate decreased cell adhesion, allowing these cells to mobilize and integrate into the elongating OFT. Because integrin signaling dynamically responds to matrix composition, these changes are expected to modulate downstream mechanotransduction pathways, thereby linking ECM remodeling to the expansion and spatial organization of the SHF niche during OFT elongation.

Because YAP/TAZ signaling is critical for maintaining SHF progenitors in a proliferative, undifferentiated state (Francou et al., 2017). The dynamic ECM remodeling observed here may therefore act as a mechanochemical switch that regulates the balance between progenitor expansion within the DPW and differentiation as cells enter the OFT.

Consistent with this model, it will be important to determine whether DPW remodeling is accompanied by changes in mechanotransduction activity, such as altered YAP/TAZ localization or integrin signaling outputs. Furthermore, the redistribution of ECM components and the apparent loss of epithelial packing raise the possibility that SHF progenitors undergo a partial epithelial-to-mesenchymal transition–like program. Future analyses examining epithelial and mesenchymal marker expression, together with quantitative assessment of progenitor cell morphology and migratory dynamics, will be necessary to determine how ECM remodeling influences SHF cell behavior and deployment during OFT elongation.

### Evolutionary conservation and developmental context of TnC function

Comparative *in silico* analysis of TnC revealed conservation of the N-terminal assembly domain, EGF-like repeats, constitutive FNIII domains, and the C-terminal fibrinogen-like globe (Chiquet-Ehrismann and Tucker, 2011) (Figure S2). Differences were restricted to the alternatively spliced FNIII cassette, a highly variable region known to vary across vertebrate species and modulate receptor interactions and matrix-binding properties (Giblin and Midwood, 2015). Thus, the divergent TnC localization in *Xenopus* is unlikely to reflect the loss of structural domains, but rather species-specific differences in splice isoform usage, ECM assembly, or mechanical context.

Alternative splicing of TnC generates multiple isoforms that differ in their interactions with extracellular matrix components and cell-surface receptors. Different splice variants display distinct adhesive properties and differential interactions with Fn1, indicating that TnC biological activity is strongly influenced by its splicing pattern (A. Ghert et al., 2001; Midwood et al., 2016). In this context, the apparent absence of several FNIII domains observed in our analysis of *Xenopus laevis* may reflect species-specific splice variants that alter TnC interactions with fibronectin and integrins, potentially contributing to the distinct ECM organization observed in the DPW.

Together, these observations suggest that although the core architecture of TnC is evolutionarily conserved, its spatial deployment within the SHF niche may be shaped by species-specific regulatory mechanisms controlling splice isoform usage and ECM organization.

### Fn1 depletion reveals its role in ECM assembly within the DPW

To further investigate the role of Fn1 in ECM organization within the DPW, we analyzed embryos depleted for Fn1 using morpholino antisense oligonucleotides. Consistent with previous studies in mammalian models, Fn1 depletion resulted in cardiac developmental defects, including shortening of the OFT, supporting a conserved role for Fn1in OFT morphogenesis (Figure 5). In addition, these embryos exhibited decreased TnC and ColI protein levels in the DPW (Figure 6). Furthermore, Fn1 depletion also appeared to alter the spatial distribution of TnC within the DPW. In control embryos, TnC is prominently detected between SHF cells across the thickness of the DPW (Figure 7). In contrast, in Fn1-depleted embryos the TnC signal is reduced, more diffusely distributed, and appears less prominent in SHF cells located farther from the endodermal interface.

This pattern suggests that Fn1 networks may be required to maintain proper ECM organization throughout the DPW, particularly in regions of the SHF spatially separated from the endoderm. Consistent with this interpretation, Fn1 fibrils are known to function as structural scaffolds that guide the incorporation and spatial organization of other ECM proteins (Sottile and Hocking, 2002). Accordingly, Fn-1 depletion resulted in decrease TnC and ColI protein levels, and altered TnC distribution, suggesting that Fn1 may act upstream in the assembly and/or stabilization of the matrix within the SHF niche, contributing to the proper positioning and retention of TnC within the matrix.

Previous studies have shown that Fn1 morpholino injection in *Xenopus* results in a strong reduction in Fn1 fibril levels during early embryogenesis, although fibril levels can partially recover at later tadpole stages as new protein accumulates (Davidson et al., 2006). Consequently, the phenotypes observed here likely reflect early defects in ECM organization rather than persistent reductions in Fn1 levels throughout all developmental stages.

Because Fn1 depletion was induced prior to the formation and deployment of cardiac progenitor populations, the resulting early disruption of ECM architecture is likely to affect the organization and behavior of cells within both the first and second heart fields. These findings support a model in which Fn1 is required to establish a permissive extracellular environment that enables the correct deployment and spatial organization of SHF progenitors during OFT elongation.

In summary, our study provides the first detailed characterization of the cellular and extracellular architecture of the DPW in amphibians, revealing a highly dynamic, stage-dependent remodeling of the SHF niche. Unlike the cohesive epithelial organization observed in mammalian models, the *Xenopus* DPW undergoes a pronounced transition into a multilayered structure. This architectural shift is closely coordinated with the spatial reorganization of the extracellular matrix, associated with the interplay between Fn1 and TnC. Our functional and *in silico* analyses suggest that Fn1 acts as a foundational scaffold required for the proper assembly and retention of TnC and ColI within the DPW. As development proceeds, we propose that species-specific alternative splicing of TnC may enable its intercellular accumulation and modulation of Fn1-mediated adhesion. This process may create a specialized mechanochemical microenvironment, potentially signaling through pathways such as YAP/TAZ, that promotes EMT-like changes in cell morphology and facilitates the deployment of SHF progenitors into the elongating outflow tract.

Although further studies will be required to directly test this model, our findings highlight that while the core molecular components of the cardiac ECM are evolutionarily conserved, their spatial organization can be highly adaptable. Understanding these species-specific differences in matrix architecture and biomechanics not only sheds light on the evolutionary plasticity of heart development but also provides important insights into how dysregulation of ECM scaffolds may contribute to congenital heart defects associated with outflow tract malformations.

## Acknowledgements

This work was supported by the Agencia Nacional de Investigación y Desarrollo (ANID) through Fondecyt de Iniciación grants No. 11240544 (CA) and No. 11220624 (PGS). IBRO RS-334417299; Centro Ciencia & Vida, FB210008 (PGS). We also thank the researchers at the Unidad de microscopía y análisis de imágenes (UMAI), Fondequip EQM190087, of CEBICEM, Universidad San Sebastián, and Light-Sheet Imaging de la Universidad Mayor (LISIUM), especially Dr. Aníbal Vargas and Dr. Luz María Fuentealba, for their assistance with the ZEISS Lightsheet 7 microscope. This equipment was funded by Fondequip EQM190087 and the LISIUM-Chile Initiative (CZI S-2022008-03).

## Author contributions

CA: conceptualization, investigation, formal analysis, methodology, and writing original draft; PGS: conceptualization, investigation, formal analysis, methodology, and writing original draft; JJ: investigation, formal analysis, and writing original draft. IS: investigation, formal analysis; CG: investigation, formal analysis; IR: investigation, formal analysis.

## Figure Legend

**Figure S1.**
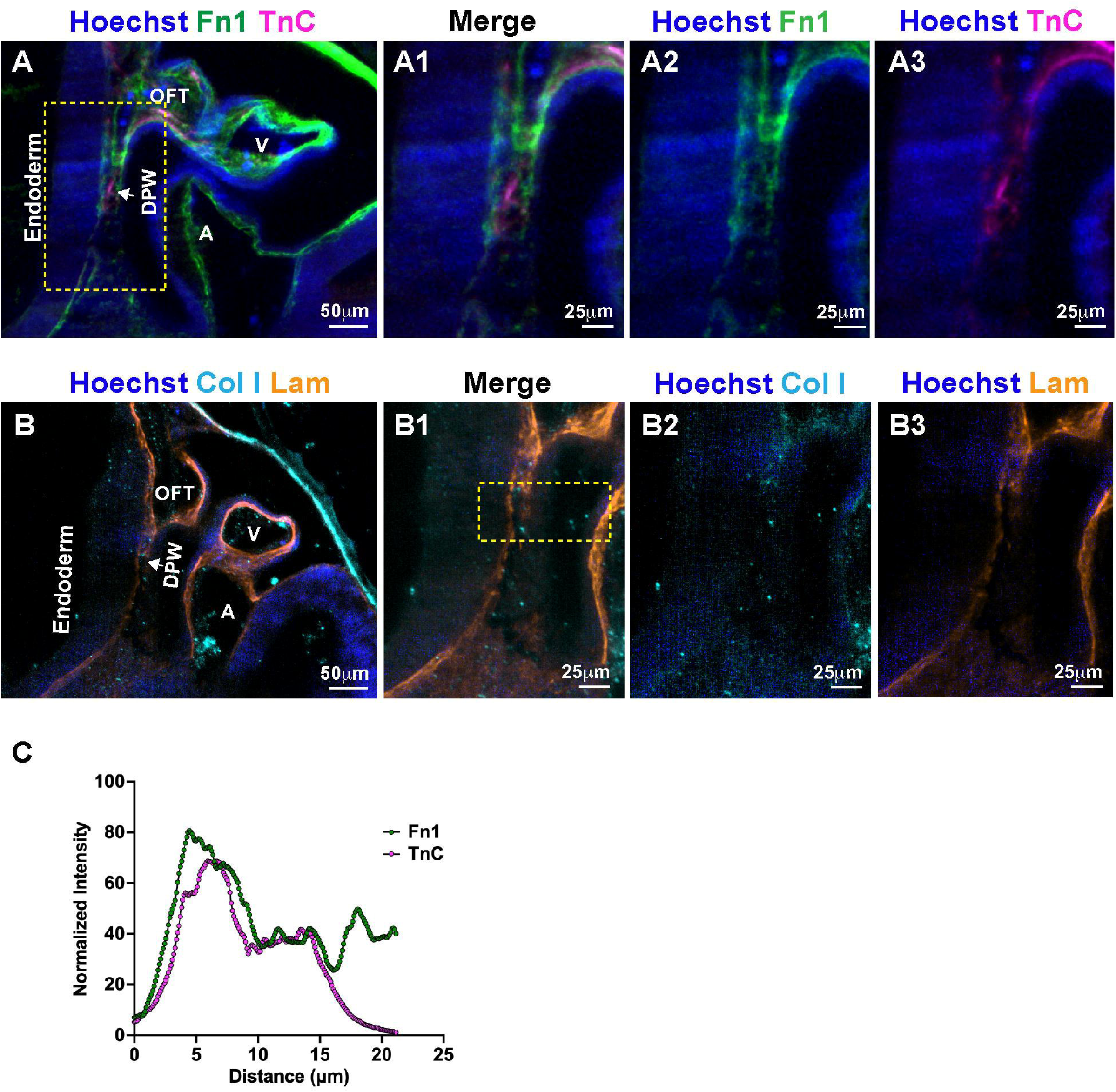
Spatial distribution and co-localization of ECM components in the developing cardiac outflow tract of *Xenopus laevis* at NF35. **A–B.** Representative sagittal sections of *Xenopus laevis* embryos showing the distribution of extracellular matrix (ECM) components in the cardiac region. Nuclei were labeled with Hoechst (blue). In (A), Fn1 (green) and TnC (magenta) are detected, while in (B), ColI (cyan) and Lam (orange) are shown. The dorsal pericardial wall (DPW), outflow tract (OFT), ventricle (V), atrium (A), and adjacent endoderm are indicated. (A1–A3, B1–B3) (C) Quantification of normalized fluorescence intensity profiles of Fn1 and TnC along a defined axis across the DPW region (as indicated in A1). Abbreviations: DPW, dorsal pericardial wall; OFT, outflow tract; V, ventricle; A, atrium

**Figure S2.**
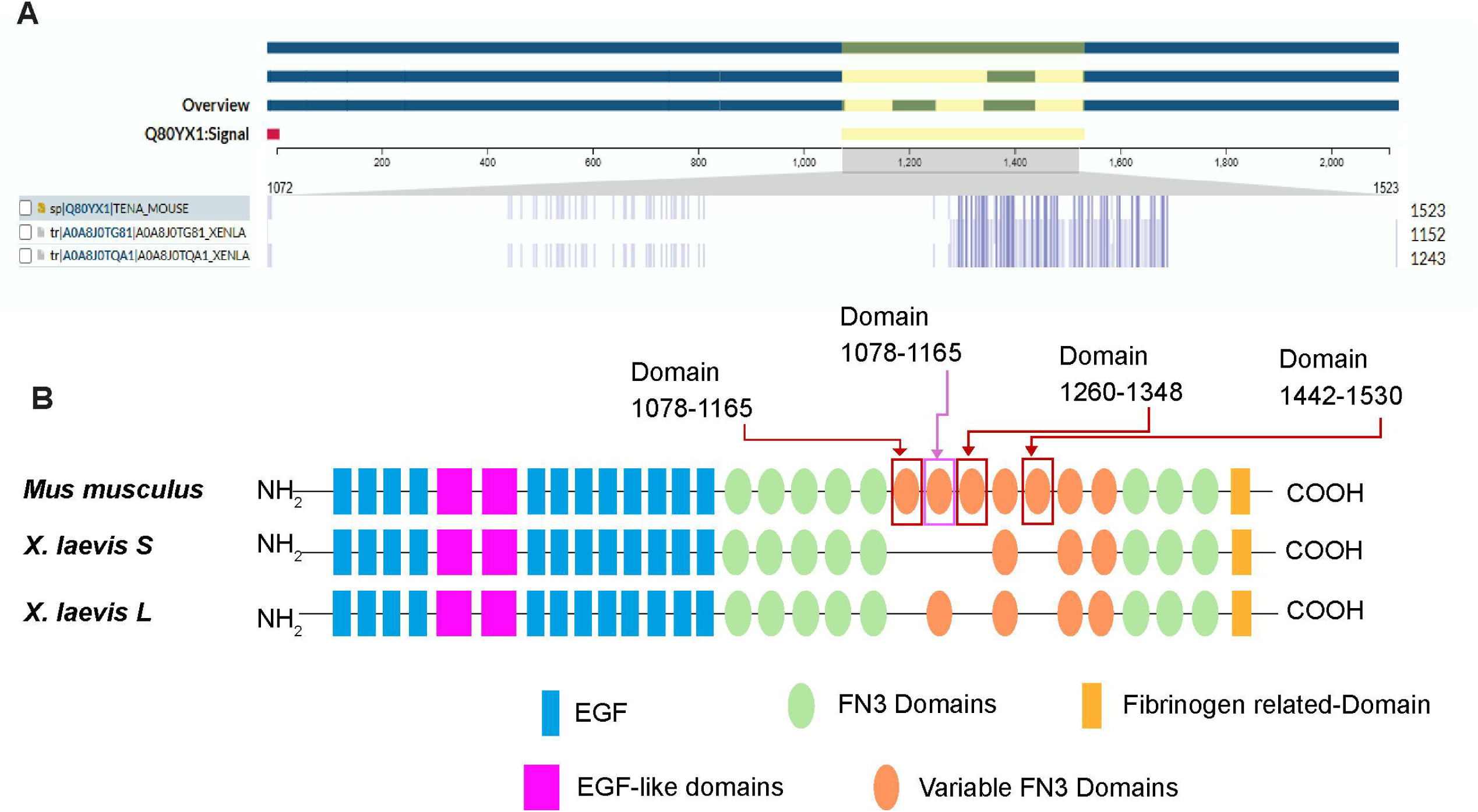
Comparative domain organization of Tenascin-C across *Mus musculus* and *Xenopus laevis*. (A) Sequence alignment overview highlighting conserved and variable regions of TnC among *Mus musculus* and *Xenopus laevis* (S and L homeologs). Conserved regions are predominantly localized within the EGF-like and FNIII domains, while variability is enriched within specific FNIII repeats, particularly in regions corresponding to alternative splicing. (B) Schematic representation of TnC domain organization. The protein consists of N-terminal EGF-like domains (blue and magenta), followed by multiple fibronectin type III (FNIII) repeats (green), including variable FNIII domains (orange), and a C-terminal fibrinogen-related domain (yellow). Comparative analysis reveals conservation of the overall domain architecture across species, with divergence in the number and arrangement of variable FNIII domains. Highlighted regions (domains 1078–1165, 1260–1348, and 1442– 1530) correspond to variable FNIII segments that may contribute to functional diversification.

**Movie 1. Cardiac morphology at NF35 in *Xenopus laevis***

3D reconstruction of the embryonic heart in Xenopus laevis at stage NF35. Nuclei are labeled with Hoechst (blue) and myocardium is visualized using MF20 (green).

**Movie 2. Cardiac morphology at NF42 in *Xenopus laevis***

3D reconstruction of the embryonic heart in Xenopus laevis at stage NF42. Nuclei are labeled with Hoechst (blue) and myocardium is visualized using MF20 (green).

## Notes

### Competing Interest Statement

The authors have declared no competing interest.

